# CCFold: rapid and accurate prediction of coiled-coil structures and application to modelling intermediate filaments

**DOI:** 10.1101/123869

**Authors:** Dmytro Guzenko, Sergei V. Strelkov

## Abstract

Accurate molecular structure of the protein dimer representing the elementary building block of intermediate filaments (IFs) is essential towards the understanding of the filament assembly, rationalizing their mechanical properties and explaining the effect of disease-related IF mutations. The dimer contains a ∼300-residue long *α*-helical coiled coil which is not assessable to either direct experimental structure determination or modelling using standard approaches. At the same time, coiled coils are well-represented in structural databases. Here we present CCFold, a generally applicable threading-based algorithm which produces coiled-coil models from protein sequence only. The algorithm is based on a statistical analysis of experimentally determined structures and can handle any hydrophobic repeat patterns in addition to the most common heptads. We demonstrate that CCFold outperforms general-purpose computational folding in terms of accuracy, while being faster by orders of magnitude. By combining the CCFold algorithm and Rosetta folding we generate representative dimer models for all IF protein classes. The source code is freely available at https://github.com/biocryst/IF

## 1 Introduction

Intermediate filaments (IFs) are an important example of a protein assembly based on *α*-helical coiled coils (CCs). IFs together with microtubules and actin filaments are key elements of the cytoskeleton in vertebrate cells. The human body contains over 70 different IF proteins, belonging to five major classes by sequence similarity and including both cytoplasmic and nuclear IFs. Inherited and sporadic mutations in IF genes were linked to numerous diseases, including muscle, heart, skin and neurological disorders (Omary, 2009), all of which are incurable at present. Currently our structural understanding of IFs is scarce, particularly when compared to that of actin and microtubules. This problem is linked to both their complexity and partial disorder. The elementary building block of all IFs is an elongated dimer with a length of ∼45 nm and a diameter of 2-3 nm. The overall structure of the dimer is defined by the formation of a CC by its conserved central ‘rod’ domain. This domain contains three CC segments denoted coil1A, coil1B and coil2 which are interconnected by short linkers L1 and L12. The rod is flanked by the intrinsically disordered, highly variable head and tail domains (Chernyatina *et al*., 2015). Following our divide-and-conquer strategy (Strelkov et *al*., 2001) multiple short fragments of the IF rod could be resolved at atomic detail using X-ray crystallography (Guzenko *et al*., 2017), while a full-length IF dimer is clearly unsuitable for crystallisation (Chernyatina *et al*., 2016).

In general, CC is a widespread structural motif in proteins involved in a multitude of functions (Lupas and Bassler, 2016). Its idealised geometry can be described with just a few parameters (Crick, 1953). The Crick parametrisation is routinely used to analyse existing CC structures (Strelkov and Burkhard, 2002) or to create theoretical models thereof (Offer *et al*., 2002; Wood *et al*., 2014). The driving force of CC formation is a regular pattern of hydrophobic amino acids that allows two or more *α*-helices to associate together, forming a common hydrophobic core. The most widespread pattern is a heptad (7-residue motif) HxxHxxx, where H indicates a hydrophobic amino acid and x stands for any amino acid. This pattern, with residue positions traditionally labelled *abcdefg*, results in a ‘canonical’ left-handed CC. Another possibility is a hendecad (11-residue repeat) HxxHxxxHxxx, which corresponds to an addition of a four-residue block (stutter) to a heptad (Lupas and Gruber, 2005). Such a pattern supports parallel packing of the chains. Two consecutive stutters after a heptad define a quindecad (15-residue repeat), promoting right-handed supercoiling. Further variation of the CC geometry is possible by insertions of three-residue blocks (stammers). In general, stammers demand a more pronounced left-handed supercoil, while stutters cause its unwinding and eventually a switch to a right-handed geometry (Lupas and Gruber, 2005).

Sequences featuring long regions with heptad patterns result in structures with a more or less uniform left-handed supercoiling and as such are the most straightforward to model (Grigoryan and DeGrado, 2011). However, sequences of many naturally occurring CCs, including those found in the IF rod, contain intermixed patterns. Experimental data reveal that such transitions can often be accommodated within a continuously *α*-helical structure, causing adaptation of the local CC parameters, so that the hydrophobic core packing is preserved (Strelkov and Burkhard, 2002). For such less regular CC structures, accurate modelling remains a challenging task. The use of Rosetta, a popular general-purpose protein folding algorithm (Leaver-Fay *et al*., 2011), for CC structures was recently described (Rämisch *et al*., 2015). Upon composing the initial fragment library from *α*-helices found in CC structures and employing an asymmetric ‘fold-and-dock’ protocol, accurate models of CCs could be obtained, including those with deviations from the heptad pattern. However, the use of Rosetta required substantial computational time (weeks) even for short CCs. For the ∼300-residue rod domain of the elementary IF dimer this is computationally prohibitive.

At the same time, the relative simplicity and the ‘linear’ nature of a parallel CC (meaning that the residues located further away in the sequence do not interact with each other) suggest that it may be amenable to reliable structure prediction. A logical option is to employ the ‘threading’ approach which can be efficient towards the *in silico* prediction of protein structure (Pieper *et al*., 2014). Indeed, there are ample experimental data for CC proteins, which can be conveniently assessed through the CC+ database (Testa *et al*., 2009). At the time of writing it contains nearly 15 thousand CCs of 11 residues or longer, 450 of which include non-canonical (*i.e*., other than heptad) patterns.

Here we describe a novel threading-based algorithm, CCFold, specifically designed for the prediction of CC structure. It is based on picking multiple CC fragments for short overlapping segments of the input sequence. For each fragment, the probability of the correct match between the sequence and structural features, such as the presence of certain residues in the core positions of the CC, is evaluated with the help of statistics obtained from the analysis of the CC+ database. The best-scoring fragments along the protein sequence are merged together through an optimization procedure. The algorithm was implemented and optimised for the parallel dimeric CCs, such as found within the IF dimer, but can also be applied to other orientations and oligomeric states. We demonstrate that CCFold clearly outperforms the existing approaches in terms of both accuracy and speed. Further, by combining the CCFold for the CC domains and general Rosetta folding we produce representative models of the elementary dimers for all main IF protein classes. Here, the availability of X-ray structures of multiple IF rod fragments allows us to further evaluate the performance of the algorithm.

## 2 System and Methods

Flowchart of the CCFold algorithm is shown in Fig. 1.

**Figure 1:**
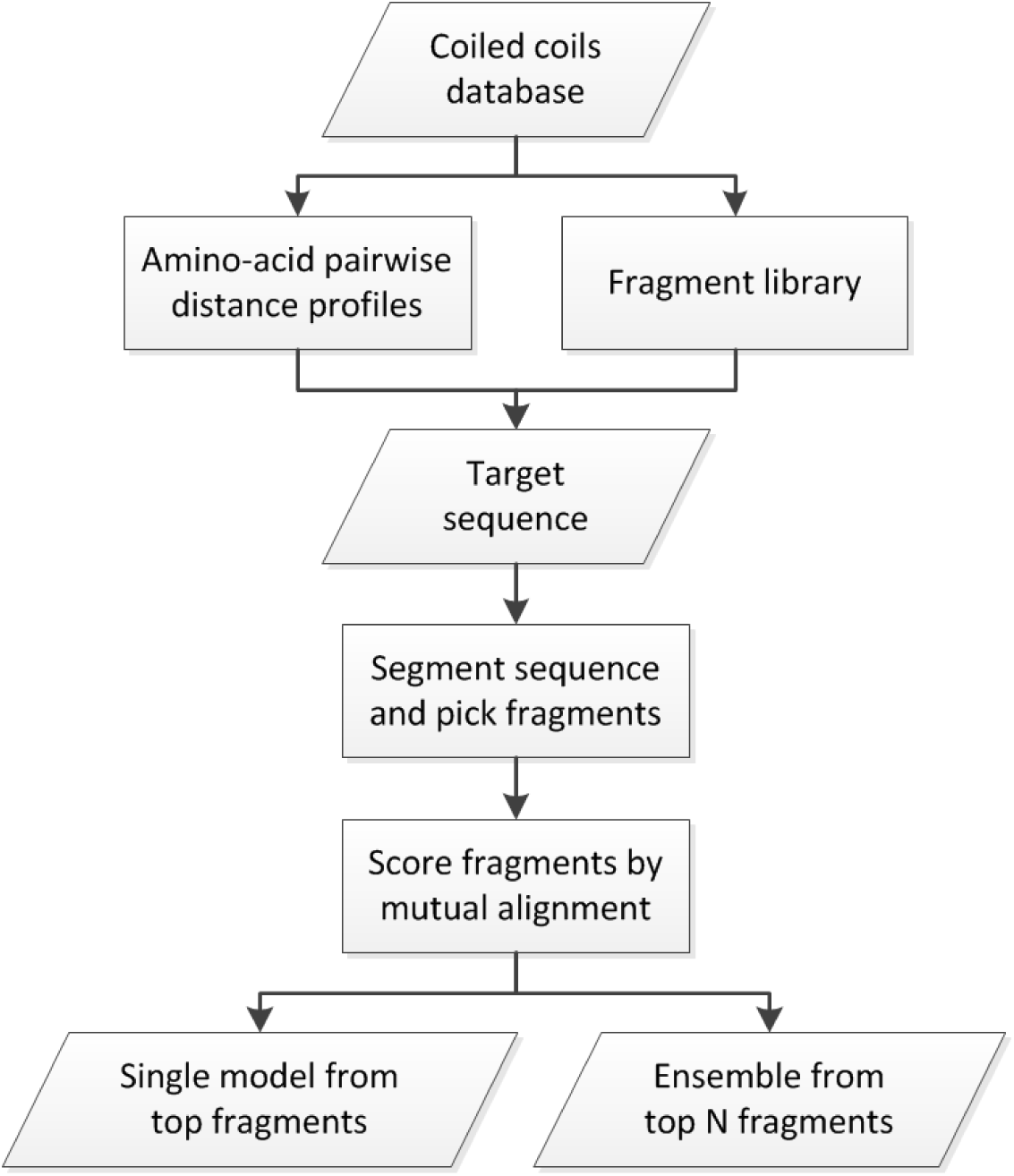
Flowchart of the CCFold algorithm.

### 2.1 Amino-acid profile of pairwise distances in parallel dimeric CCs

All parallel in-register dimeric CCs were extracted from the CC+ database and truncated to polyalanine. Chain sequences were clustered at 80% sequence identity, which resulted in 336 non-redundant CCs containing from 15 to 148 residues per chain. Statistics was collected on the distance between the C*α*-atoms of aligned residues in both chains (such as the equivalent residues in the case of parallel in-register homodimers). The number of distances measured for a particular amino acid ranged from 21 (proline) to 1899 (leucine), making a total of 11847 measurements. As seen in Fig. 2 and S1, the distributions readily reveal three major maxima, which structurally correspond to residues situated in heptad positions *a* or *d* (shortest distance), *e* or *f* (intermediate distance), and *b, c* or *f* (longest distance), respectively. At the same time, the distance distributions vary greatly across the 20 amino acids, which ultimately appears the main reason towards the high predictive power of our algorithm.

**Figure 2:**
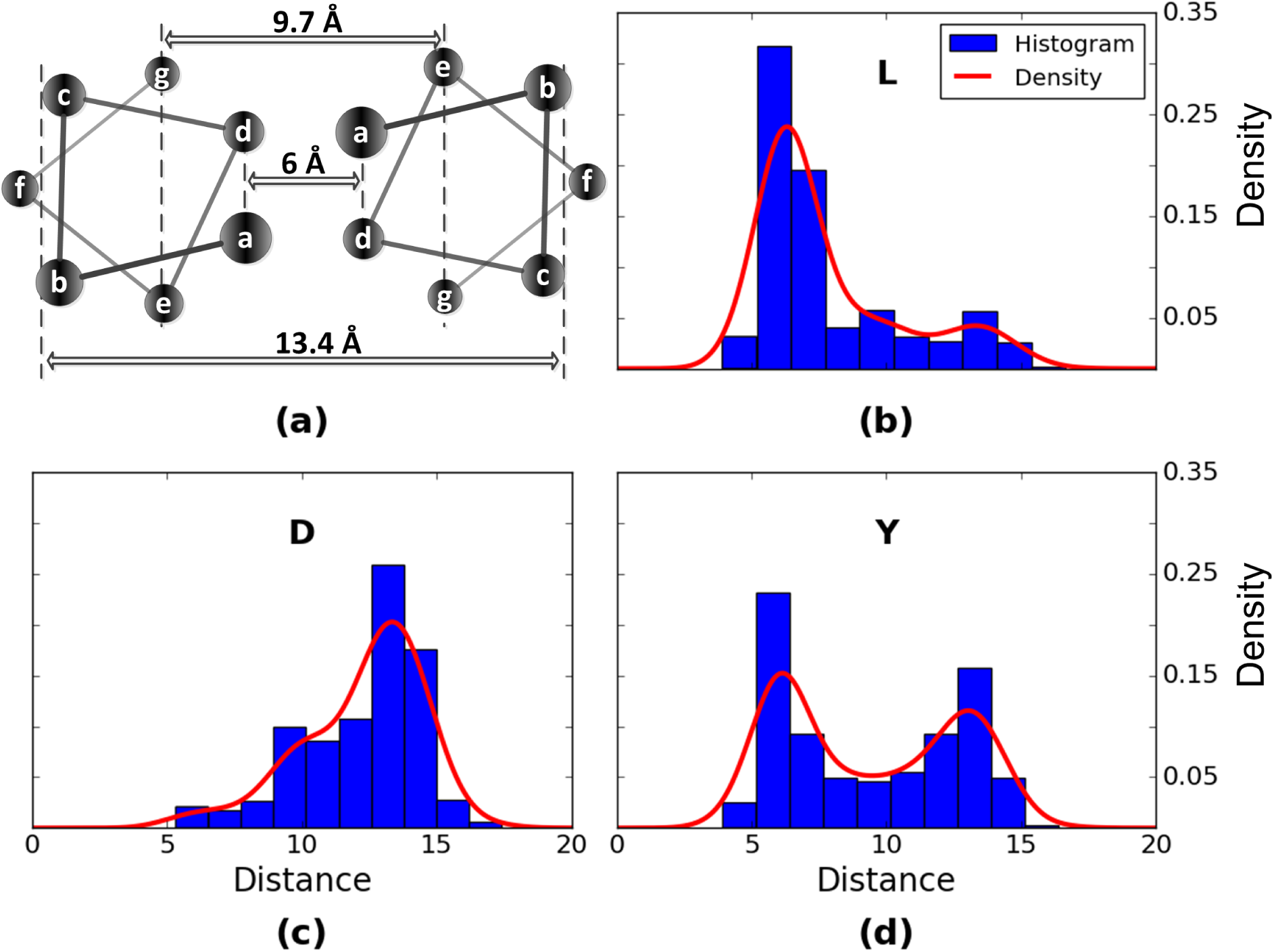
(a) Schematic diagram of an idealised CC dimer showing pairwise distances between C*α* atoms of the corresponding residues in both chains. (b,c,d) Distributions of such distances obtained from the CC+ database for valine, aspartic acid and tyrosine residues, respectively. Kernel density approximation is plotted over the original histogram.

### 2.2 CC fragment library

Next, we wanted to capture the structural diversity of experimentally determined CCs, regardless of the particular amino-acid sequences that produced such structures. To this end, a set of all possible 2x15-residue fragments of parallel two-stranded CCs was extracted from the CC+ database, truncated to polyalanine. After all-*vs*.-all coordinate root-mean-square deviation (RMSD) comparison, we selected a non-redundant subset having a pairwise RMSD between any two fragments of at least 0.2Å. This resulted in a library of 14175 dimeric CC fragments void of sequence information.

### 2.3 Delineation of CC domains

The region of the target protein sequence that is expected to form a CC structure can be predicted using standard tools (Li *et al*., 2015). In addition, the expected multiplicity of the CC (dimer, trimer, *etc*.) and the orientation of the helices (parallel or anti-parallel) must be provided. Several methods to predict the CC multiplicity exist (Vincent *et al*., 2013; Trigg *et al*., 2011). The current implementation of CCFold is focused on parallel dimeric CCs, with limited support of the anti-parallel orientation for benchmarking purposes.

## 3 The CCFold algorithm

Starting input is the amino-acid sequence to be assembled into a parallel in-register dimeric CC. The main steps of the algorithm include segmenting this sequence into a series of overlapping sequence windows, picking a pool of fragments for each window, selecting a set of fragments that optimally agree with each other in the overlapping regions, and finally merging them together.

### 3.1 Sequence segmentation

For a target sequence of length *N*, initially a full set of *N –* 14 possible overlapping sequence windows of length 15 is considered. For every window we evaluate the probability that it would form each of the fragments from our library, assuming independence between the amino acid pairs:

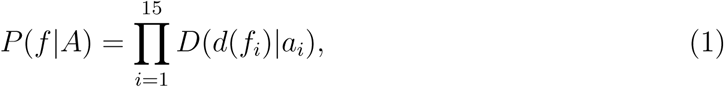
where *a_i_* is the *i*-th amino acid of the sequence window *A, d*(*f_i_*) is the distance between the aligned C*α* atoms of the *i*-th residue within the fragment *f*, and *D*(*d*(*f_i_*)*|a_i_*) is the kernel density function for amino acid *a* (Fig. 2). Importantly, the function (1) correlates well with the likeness to the true structure in terms of minimal coordinate RMSD even for non-canonical CCs. Fig. 3 demonstrates this for an experimentally determined coil2 fragment of lamin A containing a stutter (Strelkov *et al*., 2004).

**Figure 3:**
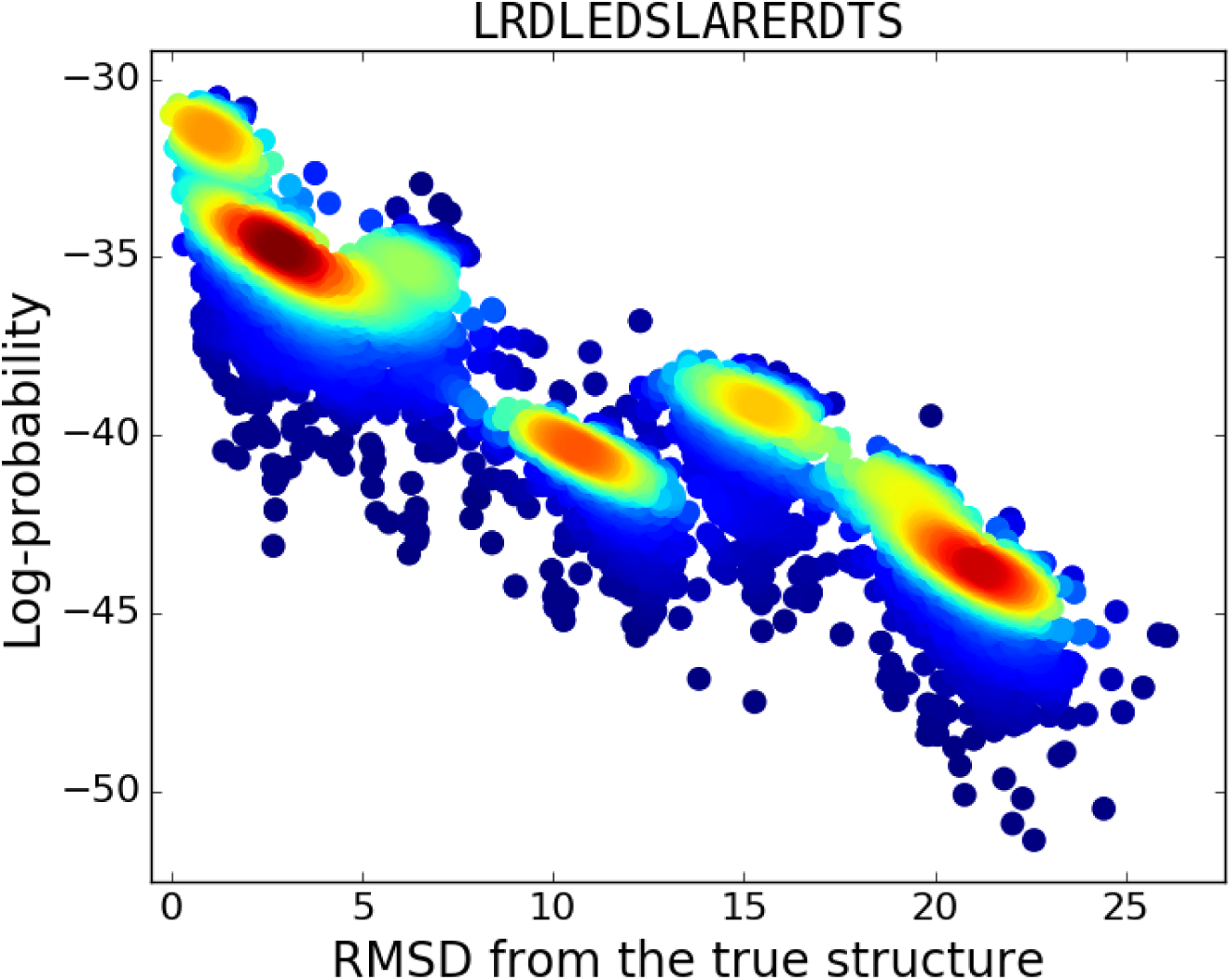
Scatter plot comparing log-probability (eq. 1) of each of 14175 database fragments for a 15-residue segment of human lamin A (residues 320 to 334) and their C*α* RMSD with respect to the corresponding part of the crystal structure (PDB entry 1X8Y). Colour indicates the data point density.

At this point, a particular set of overlapping 15-residue sequence windows covering the entire target sequence is selected. To this end, 100 top structural fragments are picked by probability (1) for every possible window. The optimal segmentation of the target sequence is given by a two-pass dynamic programming as detailed below.

Let **F***_i_* be a set of fragments for the *i*-th sequence window and let *o_i_* be a set of overlapping residues between fragment sets **F***_i_* and **F***_i_*_+1_, number of which is bounded by reasonable values *o_min_* ≤ |*o_i_*|≤ *o_max_*. The segmentation of the target sequence is defined as 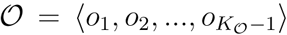, with number of resulting windows 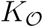 varying for different segmentations. Let 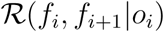 denote the RMSD of two fragments *f_i_* and *f_i_*_+1_ from consecutive fragment sets **F***_i_* and **F***_i_*_+1_ superposed by the residues in *o_i_*. Further, let 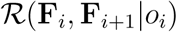 denote the average RMSD of all fragments from **F***_i_* and all fragments from **F***_i_*_+1_. The first pass of dynamic programming is employed to minimize the maximal average overlap RMSD:

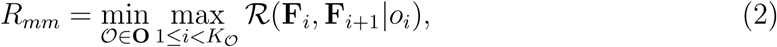
where **O** is a set of all possible overlaps. Afterwards the second pass minimizes the total alignment RMSD under condition that no single one surpasses the value determined at the first step:

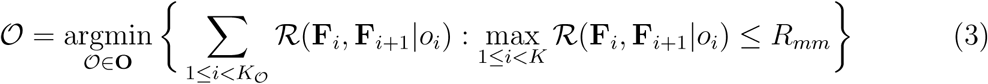

An example of a resulting sequence segmentation is shown on Fig. 4. We have found that such an approach yields more accurate models compared to just using a fixed set of overlapping fragments with constant overlap, especially in the cases when non-canonical repeats are present or the hydrophobic core assignment is ambiguous.

**Figure 4:**
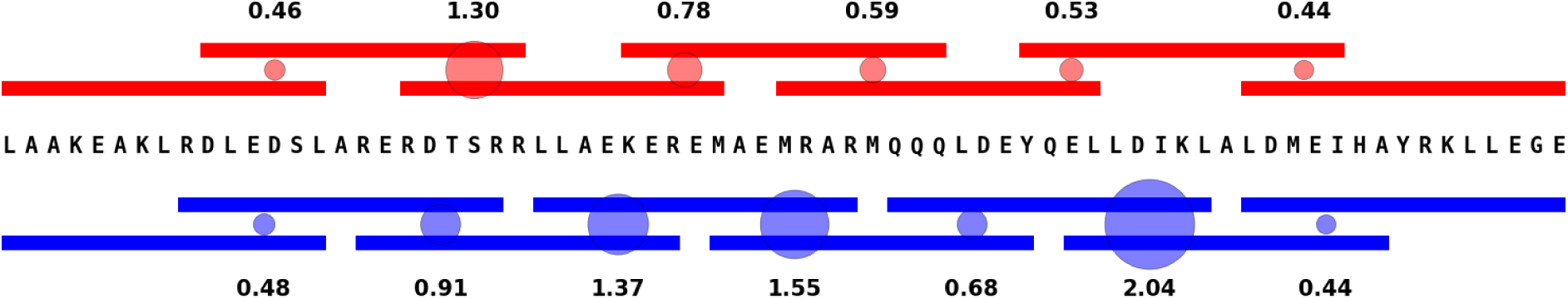
An example of sequence segmentation. Optimized set of overlapping 15-residue windows for a lamin A fragment (residues 313 to 383), shown on top. For comparison, a simple set of windows with constant overlap is shown at bottom. Average overlap RMSD (in Å) between the fragments in the windows is illustrated by circles.

### 3.2 Optimal set of overlapping structural fragments

Next, for each sequence window from the selected overlapping set we choose a large pool (200-1000) of the most probable structural fragments. Logically, the majority of such fragments in the majority of windows should represent the correct fold for a given sequence. Based on this assumption, we re-score the fragments in the pools according to how well they align with fragments selected in other windows, using sum-product belief propagation (Barber, 2012). The factor graph required for this procedure is constructed by representing each sequence window by a variable node, and every overlap between two windows by a factor node that specifies alignment RMSD between the overlapped fragments.

### 3.3 Output model generation

As the first option, the final model is constructed by simply merging the top-scoring fragments in each window. To this end the fragments are superimposed upon RMSD minimisation. Thereafter the atomic coordinates in the overlapping parts are averaged with weights decreasing towards the ends of the merged fragments. The resulting single model should be the most accurate one locally, but its overall shape (such as the bending of the CC axis in particular) is not controlled.

Alternatively, an ensemble model can be output. The advantage of such a model is that it collectively reveals the uncertainty of the structural prediction. In addition, ensemble models may be desired for some applications such as for phasing X-ray data using molecular replacement (MR). The procedure to obtain an ensemble model is as follows.

Let each sequence window be represented by any of 20 to 100 of highest-scoring fragments. This enables the solution of a ‘shortest path’ problem, *i.e*., choosing fragments that result in the lowest sum of overlap RMSDs throughout the whole structure. Let us define a *conditional* shortest path, the condition being that the path is restricted to a particular fragment *f* in a certain window **F***_j_*. With **F**\**F***_j_* = **F**_1_ ×… × **F***_j_*_–1_ × **F***_j_*_+1_ ×… × **F***_K_* we define:

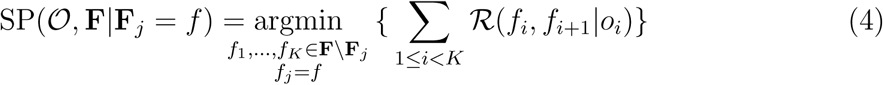

Next, we condition eq. (4) on all fragments in all windows to produce an ensemble model as follows:

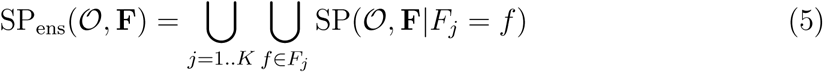

Each individual model resulting from the procedure (5) will differ from the rest in at least one 15-residue fragment. Moreover, choosing a different fragment in one window may cause selection of alternative fragments in other windows to satisfy the RMSD minimisation criterion.

Finally, the procedure (5) can be used to produce a model with a nearly straight CC axis at a small cost to the local accuracy. Let the sequence *S* = ⟨*f*_1_, *f*_2_,…, *f_K_*⟩ of chosen fragments resulting from eq. (4) be characterised by the total fragment overlap RMSD 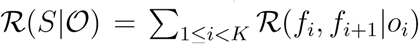, which naturally reflects overall quality of the produced model. As an indicator of straightness, we use superposition RMSD of the two chains of the full-length model 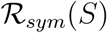. To combine these two measures, models with higher-than-median overlap RMSD are discarded from the ensemble, and the model with minimal inter-chain RMSD is selected among the rest of the ensemble:

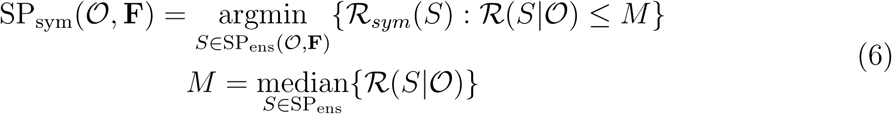

An example of output models depending on the options chosen is given in Fig. S2.

## 4 Implementation

The algorithm is implemented as a Python script. Biopython (Cock *et al*., 2009) is used to process the Protein Data Bank (PDB) files (Berman *et al*., 2000). Structure superpositions and RMSD calculations are done with PyRMSD (Gil and Guallar, 2013). The smooth density functions are estimated using the Gaussian kernel available in Scikit-learn (Pedregosa *et al*., 2011), with the bandwidth set to 1.

Program parameters are as follows:

- target=one,termini,all. Whether to produce one model, a collection of models with alternative terminal fragments, or a collection with alternative fragments in all positions.
- straighten=true,false. Whether to produce models with a straight axis.
- segmentation^·^pool (default 100). Number of fragments used to determine the optimal segmentation of the sequence.
- belief^·^propagation^·^pool (default 350). Number of top fragments in each sequence window selected for the belief propagation procedure.
- shortest^·^path^·^pool (default 50). Number of top fragments in each window selected for producing the shortest paths given the segments. Not applicable if target=one.

## 5 Modelling of the IF rod domain

The overall flow of the modelling is shown in Fig. 5.

**Figure 5:**
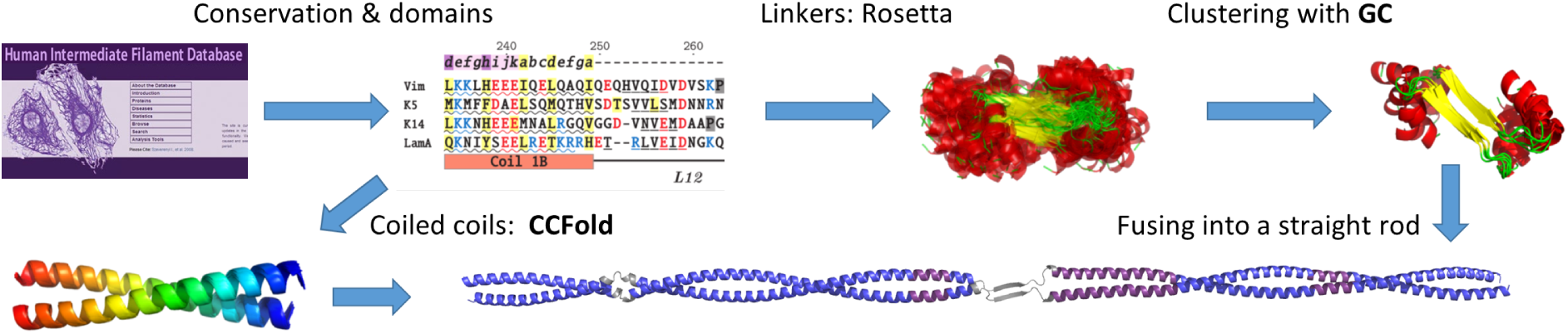
Flowchart of the IF rod domain modelling.

### 5.1 CC segments of the IF rod

IF protein sequences were retrieved from the Human Intermediate Filament Database (Szeverenyi *et al*., 2008). The CC segments (coil1A, coil1B and coil2) of each protein were defined as in Chernyatina *et al*. (2015) and fed into the CCFold algorithm. In case of heterodimers, such as for type I/II keratins, these were pairs of aligned sequences. Ensembles including 100 models with alternative N-and C-terminal fragments were produced. Straight models were preferred.

### 5.2 Linkers L1 and L12

Rosetta modelling suite (Leaver-Fay *et al*., 2011) was employed as general structure prediction method for the short linkers that interconnect the CC segments. The asymmetric fold-and-dock protocol (Rämisch *et al*., 2015) was used. This was necessary for the modelling of heterodimers such as type I/II keratins. In addition, for linker L12 a parallel *β*-hairpin structure with a two-residue offset of the two chains could be stably produced.

To this end, the sequences of the linkers were first ‘capped’ with eight-residue stretches of an *α*-helix at either end. ‘AtomPair’ distance constraints of 6Å were placed on the C*α*-atoms of hydrophobic core residues of these ‘CC caps’, to simulate the context of a dimeric rod. 5000 decoys were produced for each linker. 500 top-scoring decoys were analysed with our GC algorithm (Guzenko and Strelkov, 2017), and those forming the largest cluster were taken. Such an asymmetric procedure implied that the CC caps at either end of the linker were generally not aligned along the same axis. While the linkers are widely assumed to serve as points of flexibility of the rod domain, a ‘straight’ model is preferable as the default conformation. Accordingly, we have implemented a genetic algorithm that modifies backbone torsion angles of the linker residues which lack secondary structure in order to bring the CC domains to the same axis, as illustrated by Fig. S3. Finally, the modelled linkers were merged with the flanking CC domains using the caps.

### 5.3 Assembly of the complete rod domain

The complete rods were constructed from the obtained pools of models for the CC segments and the two linkers by superposing the overlapping parts. The side chains were then added with SCWRL4 (Krivov *et al*., 2009). Finally, the complete models were energy-minimized using REFMAC5 (Murshudov *et al*., 2011).

## 6 Results

### 6.1 CCFold algorithm validation

The CCFold algorithm enables modelling of dimeric CC structures with just the amino-acid sequence on input. The procedure is fast, allowing complete modelling of a ∼100-residue CC within a few seconds when using a regular contemporary PC. In particular, we have modelled 10 dimeric CC structures previously used as a benchmarking set for the asymmetric fold-and-dock (AFnD) protocol (Rämisch *et al*., 2015). Anti-parallel dimers were modelled with a modified version of the CCFold algorithm utilising a database of anti-parallel fragments and corresponding pairwise distance distributions. To prevent any bias from the known experimental structures, homologues of the target structure at PSIBLAST E-value of 0.05 were excluded from the database.

The models output by CCFold and the best decoys obtained by the AFnD protocol were both compared to the experimentally determined structures using two orthogonal criteria, namely the lDDT score which evaluates the local model quality (Mariani *et al*., 2013), and the TM score which assesses the similarity of the global fold (Zhang and Skolnick, 2004). Only C*α* atoms were considered. As seen from Table 1, 8 out of 10 resulting models produced by our method are superiour by both local and global similarity measures. The remaining two targets show a better result by one of the measures and worse by the other. At the same time, our algorithm is radically faster than the Rosetta-based AFnD procedure.

**Table 1:** Performance comparison of CCFold and AFnD. Better values are in bold.

| PDB ID | IDDT score |  | TM score |  |
| --- | --- | --- | --- | --- |
|  | CCFold | AFnD | CCFold | AFnD |
| 1hf9 | **0.809** | 0.750 | **0.855** | 0.668 |
| 1pl5 | **0.967** | 0.931 | **0.935** | 0.915 |
| 1r48 | **0.845** | 0.810 | **0.781** | 0.728 |
| 1t3j | **0.920** | 0.881 | **0.884** | 0.866 |
| 1uii | 0.867 | **0.910** | **0.866** | 0.850 |
| 1x8y | **0.893** | 0.811 | **0.936** | 0.857 |
| 2oqq | **0.942** | 0.941 | 0.924 | **0.930** |
| 2q6q | **0.981** | 0.963 | **0.961** | 0.923 |
| 2w6a | **0.965** | 0.943 | **0.933** | 0.925 |
| 3bas | **0.946** | 0.910 | **0.917** | 0.868 |

### 6.2 IF dimer structure

We have used the CCFold algorithm to produce representative rod domain structures for each of the main sequence homology classes of the IF family (Fig. 6). The resulting models were superposed with the available crystal structures of the rod fragments. The structural agreement is quite good, as the observed deviations are largely confined to the bending of the CC axis of the experimental structures, which are likely due to the crystal contacts they are involved in.

**Figure 6:**
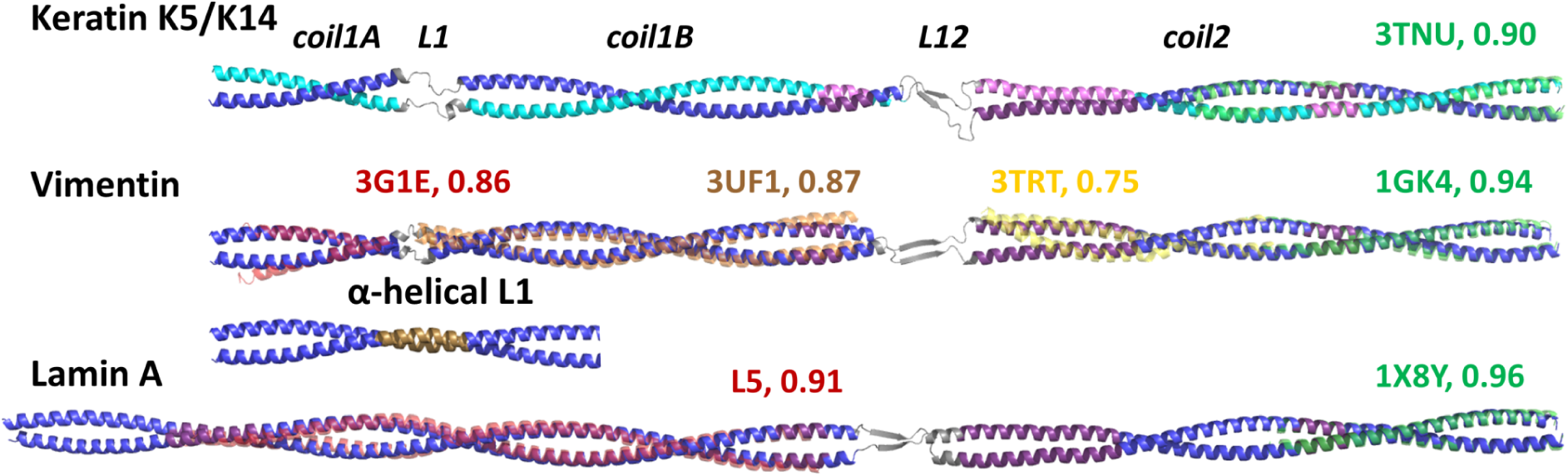
IF rod domain models. Regions with hendecad repeats are highlighted in violet. Linker regions are shown in grey. A possible fully *α*-helical conformation of linker L1 in vimentin is also shown, with 19-residue repeat highlighted in brown. The model is superposed with the available crystal structures of fragments, with PDB codes and TM scores of the superposition indicated.

Most structural features of the rod are preserved across various IF classes, which is in line with a considerable sequence conservation (Guzenko *et al*., 2017). Coil1A always appears as a relatively short CC segment with a regular left-handed structure. Coil1B also consistently reveals a left-handed structure for the most part (whereby coil1B of lamins contains an insert of 42 residues or 6 heptads compared to cytoplasmatic IF proteins), except for a single hendecad and therefore a local unwinding near its C-terminus. Finally, coil2 domain is absolutely conserved in both length and hydrophobic repeat pattern throughout all IF classes, and contains a parallel *α*-helical bundle (3 hendecad repeats) at its N-terminus, plus a single extra hendecad repeat (equivalent to a stutter) approximately two thirds into its length (Fig. 6).

In line with previous analyses (Kapinos *et al*., 2010), the linker L1 of nuclear lamins could be modelled as fully *α*-helical, so that a continuous left-handed CC is formed running from coil1A into coil1B, with a local unwinding (2 hendecads) at the linker. In cytoplasmic IF proteins, this linker was traditionally considered as a non-helical and flexible (Smith *et al*., 2002), although recent experimental data suggested a rigid structure (Aziz *et al*., 2012). Interestingly, by feeding the complete coil1A-L1-coil1B sequence of vimentin into the CCFold algorithm we could obtain a continuous CC with a single 19-residue motif at positions 135-153. While experimental verification of the latter possibility is still necessary, in Fig. 6 we present two alternative conformations for vimentin linker L1. Finally, in line with earlier sequence-based predictions (Parry and Steinert, 1999), our modelling of linker L12 yielded a short stretch of a *β*-structure in all IF types. Interestingly, the predicted parallel *β*-hairpin often reveals a two-residue shift of one chain relative to another.

We have further assessed the quality of the predicted IF rod structure by analysing its CC geometry using the program Twister (Strelkov and Burkhard, 2002). Indeed, comparison of the geometrical parameters such as CC pitch and radius for the *in silico* model and crystal structures is an efficient means to evaluate the local modelling accuracy. Fig. S4a shows the inverse of the CC pitch angle which reveals the local CC geometry (left-handed or right-handed supercoiling and its pitch), along the length of our vimentin rod model and in experimental fragments. The local unwinding of the left-handed CC at the positions of hendecad repeats, as manifested by a pronounced drop in the inverse CC pitch value, is consistently observed in both our model and crystal structures. At the same time, our model has a practically constant CC pitch (∼147 Å) and CC radius (∼4.9 Å) throughout the regions with regular heptad repeats, apparently corresponding to the average values in the fragment database, while the experimental structures reveal more local deviations (Fig. S4b). Although some of the latter may be related to both crystal contact artefacts and truncation effects, there can also be structural reasons, in particular the size of the side chains in core positions, towards ‘real’ variations of CC radius and pitch along the length of the structure. Indeed, for CC fragments crystallized as several copies in the asymmetric unit, similarities in the CC pitch profile were observed (Strelkov and Burkhard (2002), Fig. 5). At this moment, the CCFold algorithm is not able to simulate these fine details. Moreover, we note that even a small variation of the CC pitch value will naturally result in a major relative rotation of the ends of a sufficiently long CC. These global differences can be decisive in some applications, such as crystallographic phasing by MR. Particularly in the latter case we recommend to use the CCFold to generate an ensemble of models, which could ultimately be crucial for a successful phasing.

In addition, using the CCFold algorithm, we have produced dimer models of isomin and crescentin, which had previously been proposed as IF-like proteins in insects (Mencarelli *et al*., 2011) and bacteria (Ausmees *et al*., 2003), respectively. In line with previous analyses (Herrmann and Strelkov, 2011) pointing to conservation of several sequence features, the isomin dimer model revealed a CC structure resembling that of the nuclear lamins (Fig. S5a). This included a short linker corresponding to L1 as well as a parallel *α*-helical bundle near the beginning of coil2. However, our new modelling suggested a possibility of a fully helical structure for the linker L12 in isomin. Also the coil2 of isomin contains an insertion of 2 residues in the place of a traditional stutter in IF proteins, resulting in an additional short linker. In contrast, our modelling of crescentin yielded a continuous CC structure without any linkers, predominantly based on regular heptads with only two instances of stutters/hendecads (Fig. S5b). We conclude that crescentin dimer bears little resemblance to the segmented IF rod domain.

### 6.3 Application to molecular replacement

By serendipity, further evidence on the quality of CC models obtained by the CCFold algorithm could be obtained. In the past, we have collected X-ray diffraction data for crystals of a lamin A fragment (residues 65-222 corresponding to linker L1 and coil1B; Chernyatina *et al*., in preparation). All previous attempts to phase the data by MR failed, despite a large number of search models tested. Indeed, the MR procedure for CC structures is known to be challenging in general and highly sensitive to the quality of the search model (Guzenko *et al*., 2017). However, an MR search using a model of this fragment produced by the CCFold algorithm was successful. The refined crystal structure (fragment L5 on Fig. 6) shows only minor deviations from the modelled structure and in particular supports a fully *α*-helical conformation for the linker L1.

## 7 Discussion

Existing methods to predict CC domains almost exclusively employ heptad repeat as the main paradigm to define a CC (Li *et al*., 2015). Correspondingly, such algorithms tend to assign lower CC scores to regions with other repeat patterns, which are only seen as ‘discontinuities’ in the heptad periodicity. With time, however, decads, hendecads, quindecads, etc. were determined experimentally to be consistent with a continuous CC structure (Lupas and Gruber, 2005). With this in mind, here we do not attempt to route the modelling of a particular protein sequence towards this or that type of CC periodicity, but pose a more general question: can this sequence be folded into a continuous CC with a plausible hydrophobic core? To answer this question we base ourselves on threading through all available experimental CC structures.

By threading fragments of CCs rather than single helices we eliminate the necessity of docking the helices together, as in the fold-and-dock approach (Rämisch *et al*., 2015). Moreover, *α*-helices forming a CC exhibit less structural variation than a combination of independent helices (Grigoryan and DeGrado, 2011). Together with fragment picking according to the amino-acid sequence and re-scoring based on mutual alignment of the fragments, this approach drastically reduces the conformational search space.

It should be noted that the starting hypothesis of a CCFold run on a given sequence is that this sequence indeed forms a dimeric parallel CC. Thus the algorithm would attempt to construct a continuous CC even in cases when this hypothesis is wrong. In order to see how the CCFold algorithm deals with this situation, we tried to deliberately feed in amino-acid sequences not forming a CC. The models output in these cases include clear structural abnormalities, for instance a deviation of the *α*-helices from the optimal geometry (3.6 residues per turn and the canonical hydrogen bonding pattern of main-chain atoms). Thus, beyond its initial purpose as a *de novo* structural modelling tool, our algorithm can be useful as a predictor of CC domains. At the same time, in borderline cases the CC structure produced by our algorithm may still need experimental verification, such as for the alternative model of vimentin rod with fully *α*-helical linker L1 (Fig. 6). The predictive power of CCFold directly relates to the quality of the underlying structural database and inclusion of various possible types of CC geometry. Thus the accuracy of our predictions should further increase with time, as more CC structures are determined experimentally.

The CCFold algorithm brings the modelling of the IF dimer structure to a new level. Past attempts included choosing some starting conformation, based either on the available crystal structures of protein fragments or some rather crude structural assumptions, followed by molecular dynamics (MD) (Chou and Buehler, 2012; Bray *et al*., 2015). The problem here is that, given the size of the IF dimer and current computational capacities, such MD simulations could never guarantee a convergence to a correct fully energy-minimised structure. Typically, only a locally optimized structure could be obtained, which was still principally defined by the starting conformation.

Here we have presented an accurate molecular structure of the conserved rod domain, the ‘signature’ feature of the IF protein class. This directly provides for the modelling of the full IF dimer, since the variable, intrinsically disordered N- and C-terminal domains can readily be simulated using standard Rosetta tools (Chernyatina *et al*., 2015). The knowledge of the dimer structure is indispensable towards the understanding of the process of IF assembly. Indeed, experimental constraints for the higher-level association of dimers are available, including chemical crosslinks in particular, as well as a more detailed information on the association of dimers into tetramers obtained through crystallography and site-directed spin labelling (Chernyatina *et al*., 2016). New improvements in cryoEM technique, as recently applied to nuclear lamins (Turgay *et al*., 2017), also suggest that locating individual dimers in tomographic reconstructions is within reach. Altogether, these advances suggest that building up a complete 3D atomic model of an IF may be achieved in the near future.

